# Elucidation of putative key genes involved in the regulation of triple negative breast cancer development and progression

**DOI:** 10.64898/2026.04.15.718835

**Authors:** Alok Kumar, Gauri Shakar Upadhyay, Mohammad Kashif, Md. Zubbair Malik, Naidu Subbarao, Maitreyi S Rajala

**Affiliations:** School of Biotechnology, Jawaharlal Nehru University, New Delhi, India; School of Computational and Integrative Sciences, Jawaharlal Nehru University, New Delhi, India

**Keywords:** TNBC, non-TNBC, Systems biology, KEGG analysis, Expression profiling, QPCR

## Abstract

The molecular basis of triple-negative breast cancer (TNBC), a highly aggressive and therapy-resistant subtype of breast cancer, is poorly understood. This study aims to identify key genes and pathways involved in TNBC development and progression using a systems biology approach followed by experimental validation. Here, two transcriptome microarray datasets from the GEO database were analysed using the R package LIMMA to detect differentially expressed genes (DEGs) in TNBC tumors. Gene Ontology (GO) and Kyoto Encyclopaedia of Genes and Genomes (KEGG) enrichment analyses using the DAVID database were performed to identify DEGs’ regulated biological functions and pathways. Further, a protein–protein interaction (PPI) network was constructed using the STRING online database, and the topological properties were determined using MCODE and Cytohubba plug-ins. The expression and the prognostic value of the hub genes were validated using the Cancer Genome Atlas (TCGA) survival analysis. We found 727 DEGs, of which 473 were downregulated and 254 were upregulated in TNBC *vs*. non-TNBC samples. The GO and KEGG analyses indicated that the DEGs were mainly related to cell adhesion, tumorigenesis, and cellular immunity. The PPI network had shown six hub genes, namely CCND1, CDH1, ESR1, FN1, IL6, and PPARG, as the top key regulators. All these genes were validated by quantitative real-time PCR in the TNBC cell line using non-TNBC cell line as a calibrator, and the obtained results were in accordance with the bioinformatics data. This information may contribute to understanding the various molecular mechanisms that drive the development and progression of TNBC tumors.

## Introduction

Breast cancer is one of the most prevalent cancers and is a major cause of cancer-related deaths in women worldwide [1]. Based on the presence of human epidermal growth factor receptor 2 (HER2), progesterone receptor (PR), estrogen receptor (ER), and ki67, breast cancer is grouped into four intrinsic molecular subtypes clinically. Triple negative breast cancer (TNBC) is a subtype characterized by the absence of HER2 amplification and lack of PR and ER expression [1]. It represents 10% to 20% of all breast cancer cases with a high morbidity and mortality [6]. TNBC patients have the worst prognosis, partly due to their inability to respond to endocrine therapy or HER2 targeted therapy [2]. It is a more aggressive and invasive type compared to other molecular subtypes of breast cancer [1–3]. Given that no specific therapy exists for TNBC tumors besides chemotherapy, immunotherapy and surgery [7-9], there is a need to identify novel therapeutic targets and to enhance our understanding of TNBC development and progression in order to develop effective treatment options.

During the past few decades, the molecular heterogeneity and clinical condition of breast cancer have been widely recognized. The development and widespread application of high-throughput technologies, including microarray and RNA sequencing, have provided a novel understanding of the molecular complexity of this disease [10-12]. Several studies have reported the expression of genes such as EGFR, KRT17, KRT14 and KRT5 in TNBC cells that are usually a characteristic of normal myoepithelial/basal cells [7, 13]. Other molecular features of TNBC include, deregulated cell cycle, inactivation of BRCA1 and BRCA2, aberrant activation of proliferative signal pathways and a high frequency of mutations in tumor suppressor genes [14].

In the current study, differentially expressed genes (DEGs) from two TNBC datasets were utilized to construct a protein-protein interaction (PPI) network. This analysis aimed to explore the fundamental topology of the network and identify novel key regulators in TNBC tumorigenesis. Gene enrichment details of the DEGs were also identified. Genes that were already identified as therapeutic biomarkers were validated through TCGA data.

## Materials and methods

### Data retrieval

TNBC datasets were retrieved from GEO database (https://www.ncbi.nlm.nih.gov/geo/, accessed on 20 December 2022). Two human transcriptomic datasets, GSE27447 (n=12) and GSE39004 (n=73), both TNBC and non-TNBC were taken for the studyGSE27447 dataset consists of 5 TNBC and 7 non-TNBC human tissue samples, while GSE39004 datasets consists of 30 TNBC and 47 non-TNBC human tissue samples.

### Data pre-processing and identification of DEGs

Patient datasets were analysed using GEO-2R (www.ncbi.nlm.nih.gov/geo/geo2r/). Two groups were defined as non-TNBC and TNBC; p-value was adjusted to Benjamini & Hochberg false discovery rate method with significance level of 0.05 and auto-detect log transformation was performed group wise. The tool compared the TNBC group with non-TNBC to identify DEGs using the R LIMMA package; p-value < 0.05. and |log_2_FC| >1 was used as a threshold as significant DEGs for each dataset. The DEGs were selected by a Bioconductor software package called LIMMA in R program language using linear models from the normalized data of TNBC and non-TNBC tissues. Expression values of DEGs with Fold_2_ Change > |1| for upregulated and down regulated genes of each dataset were selected for further analysis. The DEGs were visualised in a volcano plot using the R package ggplot2. The BRCW computing website (http://jura.wi.mit.edu/bioc/tools/compare.php) was used to choose unique DEGs that were shared by at least two gene expression profile datasets. As a result, DEG selection was more precise, and the possibility of biased data compilation was minimized.

### Gene ontology analysis

The DEGs functional interpretation was evaluated and visualized in Database for Annotation, Visualization and Integration Discovery (DAVID; version 6.7, accessed on 7 January 2022 [15] for the molecular function, biological process and cellular component. For the metabolic pathway enrichment study, the KEGG [16] tool was used. The adjusted p-value < 0.05 cut-off score was taken into consideration for statistical significance.

### Protein-protein (PPI) network construction and analysis

The STRING database was used to extract interconnected genes to create a network of PPI [17]. To visualize PPI, the tool Cytoscape version 3.8.2 [18] was used. While, Cytohubba [19] and Network Analyzer plugin of Cytoscape was used to calculate the topological properties of PPI network. In addition, topological properties such as degree, betweenness centrality and bottleneck were used for identification of key genes, of which 20 were found to be the significant regulators. In order to investigate the essential behaviour of the network, the following properties were analysed.

#### Degree(k)

In the process of network analysis, the total number of links is established by a node and is indicated by the degree *k*. This is also used to measure the local significance of a node in the network regulation process. In the graph represented by *G = (N, E), N* denotes the nodes while the edges are denoted by E. The degree of *i*^th^ node (*k*_*i*_) is expressed as 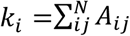, where the adjacent matrix elements of the graph are denoted by *A*_*ij*_.

#### Betweenness

The ***C***_***B***_ or betweenness centrality measures a node’s influence quantifying how often it acts as a link along the shortest-path between all pairs through nodes *i* to *j*. So, it is the ability of a node to extract benefit from the flow of information throughout the network [20] and its ability to control the signal processing over the other nodes within the network [21]. If *d*_*ij*_(*v*) represents the number of geodesic paths from one node *i* to another node *j* passing through the node *v*, then *C*_*B*_(*v*) of node *v* can be derived by the equation 2.

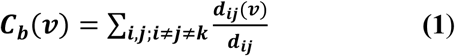

The normalized betweenness centrality is summarized in the equation 3, in which *M* represents the number of node pairs, excluding *v*.

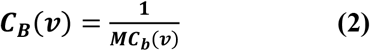

#### Bottle Neck (B_N_)

The high betweenness nodes are the bottleneck, it can be estimated using centrality betweenness, which seems to be a measurement of the centrality of a node in a network, and equal to the number of shortest paths, Let Dn be the shortest node-rooted path tree,

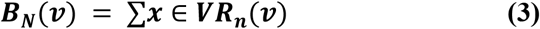

where *R*_*n*_(*v*) = 1 if more than 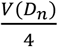 paths from node *n* to other nodes in *D*_*n*_ meet at the vertex *v*; otherwise *R*_*n*_(*v*) = 0

### Genetic alteration and validation of key genes expression paradigm

The GEPIA data tool was used to see whether the expression of key genes was linked to the survival of breast cancer patients from the TCGA data. The breast cancer patients were divided into two classes based on median gene expression values. Their overall survivals (OSs) were evaluated using the Kaplan–Meier approach with a log-rank test for obtaining the survival plot of the key gene [23]. The key genes were validated with a box plot, analysed according to the pathological stage and transcript per million. The p < 0.05 was considered statistically significant.

### Validation of key gene expression in TNBC patients using UALCAN database

The expression of key genes was further validated using the UALCAN web portal (https://ualcan.path.uab.edu/index.html), which derives transcriptome data from TCGA and allows analysis of gene expression across various cancer subtypes [46]. In this study, the expression levels of the identified hub genes were assessed in samples of breast invasive carcinoma (BRCA). Gene expression analysis was conducted by utilizing the “TCGA Gene Analysis” module and inputting the gene symbols one at a time. The transcript expression levels were compared across normal breast tissues and various molecular subtypes of breast cancer, which include luminal, HER2-positive, and TNBC samples. The gene expression values were reported as transcripts per million (TPM). The boxplots generated by the platform were used to visualize the relative expression levels of the hub genes.

### Quantitative real time PCR

To experimentally validate the expression patterns of the identified hub genes, quantitative real-time PCR (qRT-PCR) analysis was carried out utilizing two breast cancer cell lines that represent distinct molecular subtypes: MCF7 and MDA-MB-231. MCF7 cells were employed as a model for luminal breast cancer (non-TNBC), while MDA-MB-231 cells acted as a representative model for TNBC. Total RNA was isolated from MDA-MB-231 and MCF-7 cells cultured in DMEM with 10% FBS by Trizol (Thermo Fisher) method. A fraction of RNA was reverse transcribed and 100ng of cDNA was subjected to real time PCR using specific primers (**CCND1**-FP: 5’TCTACACCGACAACTCCATCCG 3’, RP: 5’ TCTGGCATTTTGGAGAGGAAGTG 3’; **ESR1**-FP: 5’AGACAGCCACTCACCTCTTCAG 3’, RP: 5’TTCTGCCAGTG CCTCTTTGCTG 3’; **CDH1**-FP: 5’GCCTCCTGA AAAAGAGAGTGGAAG 3’, RP: 5’TGG CAGTGTCTCTCCAAATCCG 3’; **FN1**-FP: 5’ACAACACCGA GGTGACTGAGAC 3’, RP: 5’GGACACAACGATGCTTCCTGAG 3’; **PPARG**-FP: 5’AGCCTGCGAAAGCCTTTTG GTG 3’, RP: 5’GGCTTCACATTCAGCAAACCTGG 3’; **IL-6-**FP: 5’AGACAGCCACTC ACCTCTTCAG 3’, RP: 5’TTCTGCCAGTGCCTCTTTGCTG 3’; **GAPDH**-FP: 5’TGCAC CACCAACTGCTTAGC 3’, RP: 5’GGCATGGACTGTGGTCATGAG 3’) to the identified key genes by the SYBR green method. Primer sequences were taken from published papers from Origene website (www.origene.com). Each sample was run in technical triplicates and the mean Ct value was taken for each set of reactions. The relative expression levels of hub genes were quantified using the 2^-ΔΔCt^ method, with the expression in MCF7 cells serving as the reference baseline.

## Results

### Identification of DEGs

A total number of 22193 and 11219 annotated transcripts obtained from the datasets GSE27447 and GSE39004_GPL6244 respectively was taken for evaluation. A total of 368 downregulated and 343 upregulated genes were selected in the GSE27447 dataset on the basis of selection criteria (*p*-value < 0.05 and [log2FC] > +1, [log2FC] < −1) in TNBC and compared to non-TNBC samples. Subsequently, 473 downregulated and 254 upregulated genes were identified in the GSE39004_GPL6244 dataset. The distribution of the DEGs in each dataset is shown in **Fig. 1A-B**. A comparison of the complete genes and top 100 genes expression profile of TNBC and non-TNBC patient samples was demonstrated in a graded manner by heatmap construction (**Fig. 1C**). Furthermore, to classify unique upregulated and downregulated genes of both the datasets, bioinformatics and the research computing online tool was used (http://barc.wi.mit.edu/tools/). From both the datasets, 816 most significant unique upregulated and 992 unique downregulated genes were obtained after assessment.

**Fig. 1:**
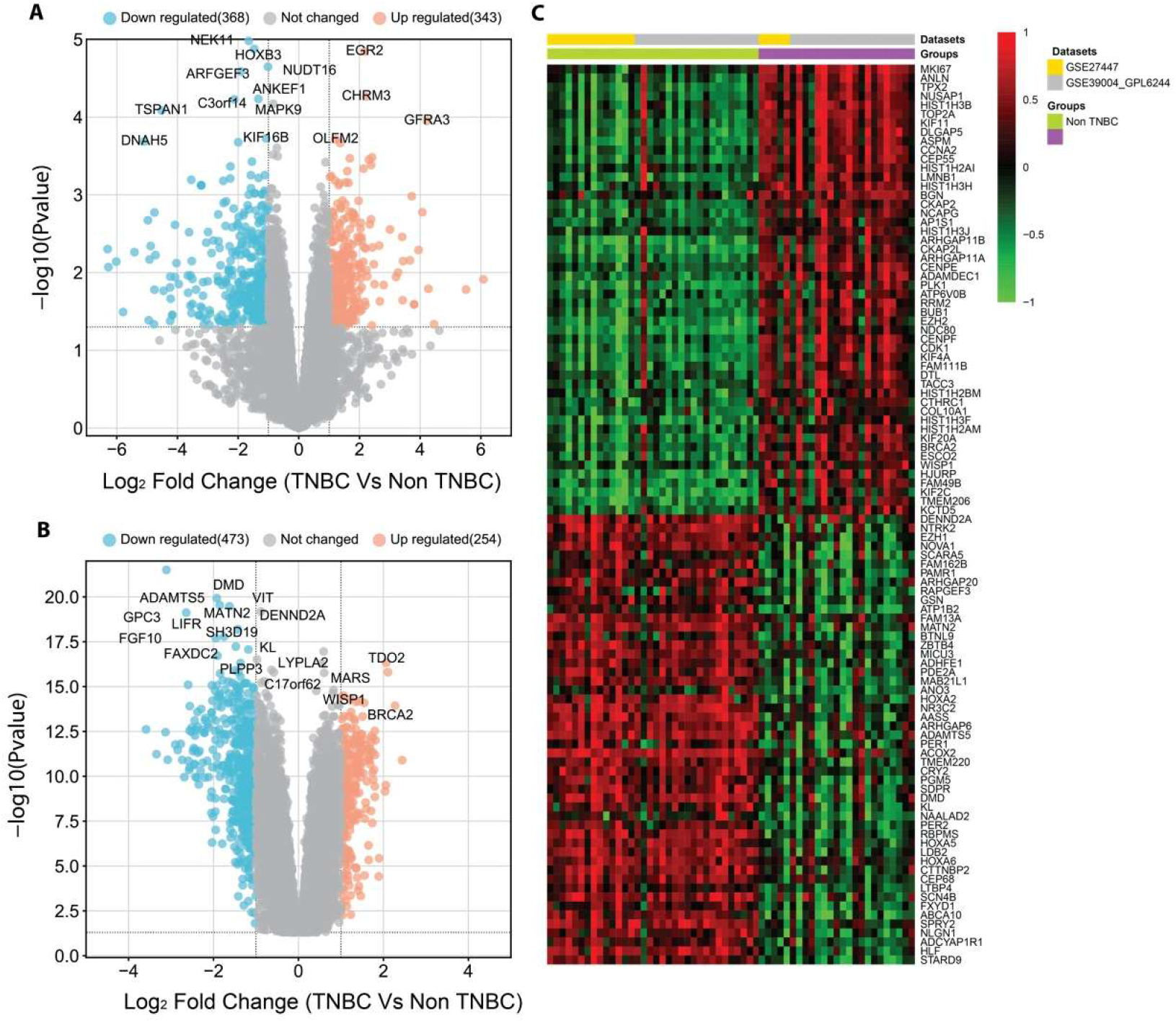
Identification of DEGs in two TNBC microarray datasets from GEO. The orange and cyan colored points denote the up and downregulated DEGs respectively. All the gray-colored points denote non-significant genes. The x and y axes represent the log2FC and −log10 p-value, respectively **(A)** Volcano plot distribution highlighting DEGs for GSE27447 between TNBC and non-TNBC samples. The orange and cyan colored points denote the up (343) and downregulated (368) DEGs. (**B**) Volcano plot distribution highlighting DEGs for GSE39004 between TNBC and non-TNBC samples. The orange and cyan colored points denote the up (254) and downregulated (473) DEGs. **(C)** The heatmap denotes the gene expression of top-100 gens.

### Gene ontology and pathways of DEGs in TNBC

Gene ontology enrichment analysis was performed to explore the biological functions, cellular component, molecular function and KEGG pathways of the identified DEGs in TNBC. The biological process category analysis had demonstrated the genes that are involved in immune response, were significantly downregulated (**Fig. 2A**). Under the cellular component, membrane and cytoskeletal elements were significantly downregulated (**Fig. 2B**). While under the molecular function category, downregulated genes were identified in association with cellular and immunological signalling (**Fig. 2C**). According to KEGG pathway enrichment analysis, the down regulated genes were significantly enriched in pathways associated with cancer, infections and diseases (**Fig. 2D**).

**Fig. 2:**
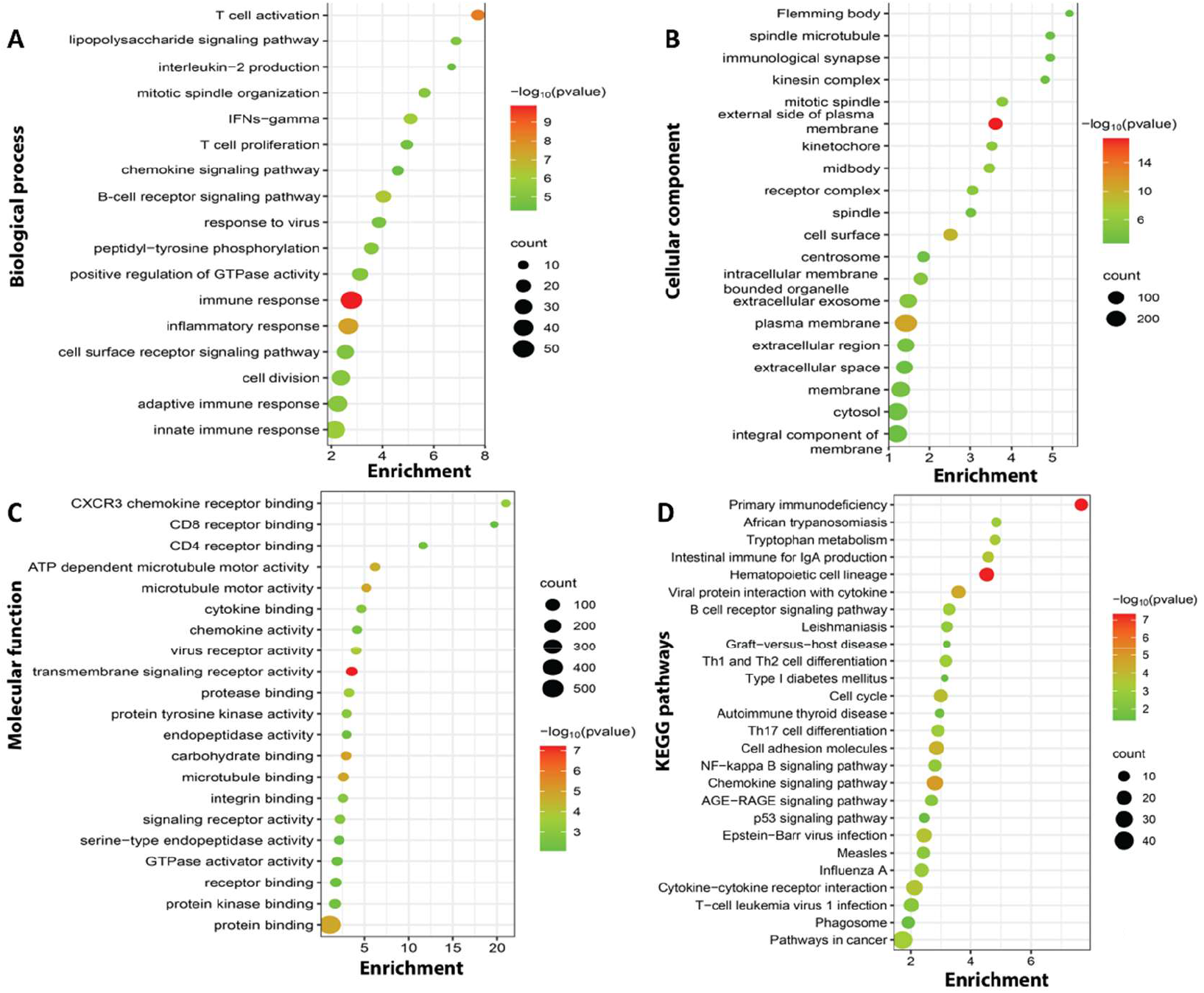
Gene enrichment analyses of down regulated DEGs in TNBC. **(A)** Significant biological processes (BP). (**B**) Enriched cellular component (CC). **(C)** Enriched molecular function (MF). (**D**) Significant enriched KEGG pathway (KP) of the down regulated genes in TNBC.

Genes that were identified to be upregulated under biological process category were associated mainly with signal transduction, cell adhesion, migration, and cellular response to external factors (**Fig. 3A**). Under the cellular component category, the up regulated genes were in association with non-nuclear components of the cell (**Fig. 3B**). For the molecular functions category, up regulated genes in TNBC were enriched in various metabolic pathways (**Fig. 3C and 3D**).

**Fig. 3:**
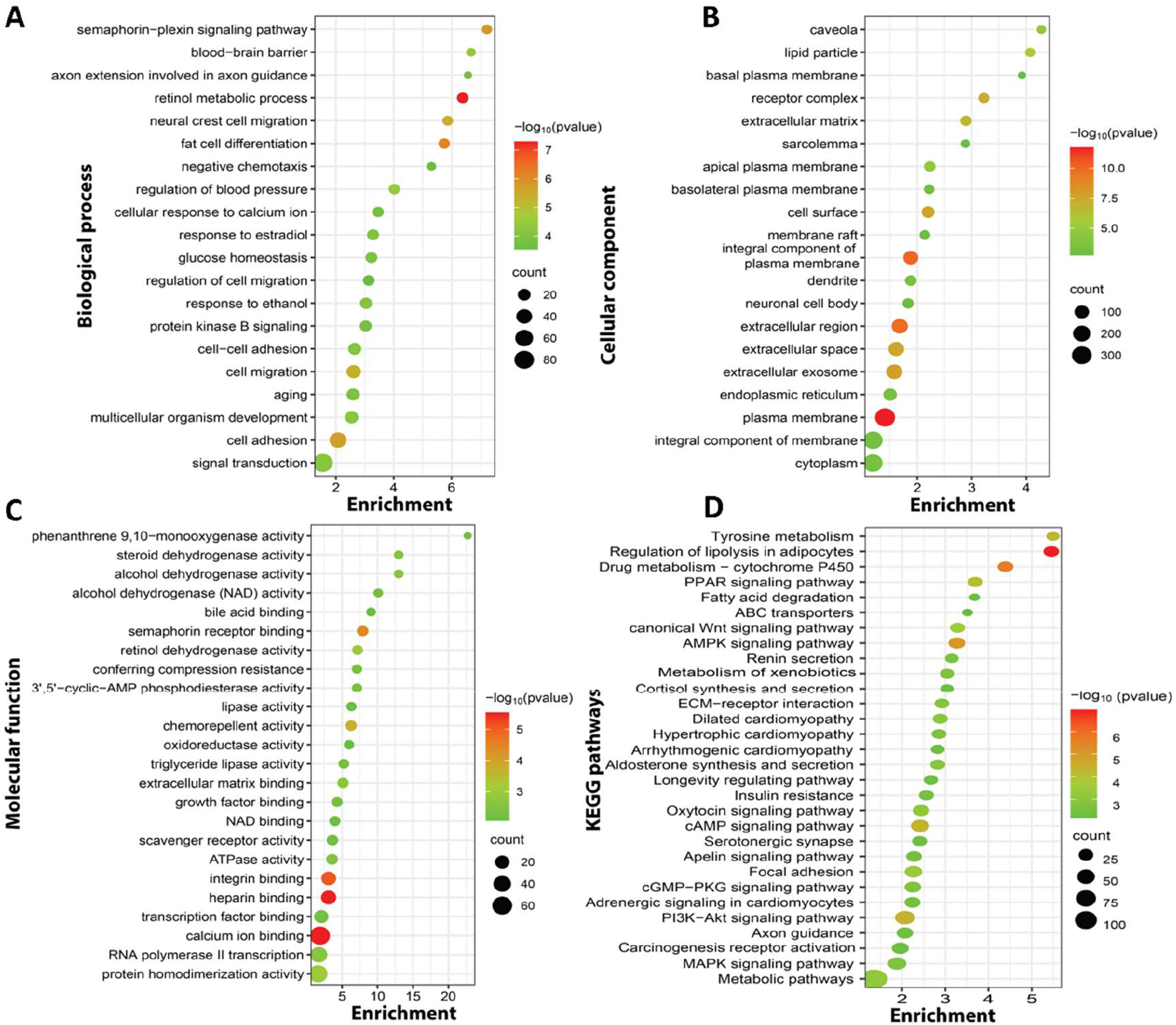
Gene enrichment analyses of up-regulated DEGs in TNBC. (**A**) Significant biological processes (BP). (**B**) Enriched cellular component (CC). (**C**) Enriched molecular function (MF). (**D**) Significant enriched KEGG pathway (KP) of the up-regulated genes in TNBC.

### TNBC PPI construction and identification of key genes

A total of 1810 DEGs were submitted to STRING for the PPI construction and analysis of the network which had shown that it has 1608 interacting nodes and 15524 edges (**Fig. 4A**). Further analysis that was executed using Cytoscape had shown 20 top hub genes (genes with highest degrees) namely IL6, PTPRC, FN1, CDH1, JUN, ESR1, CCND1, MKI67, ERBB2, CDK1, CCNB1, CCNA2, EZH2, MMP9, PPARG, CD34, CCL2, CD19, KIF11 and CXCR4 as per the decreasing order of degree values (**Fig. 4B**). The top 20 higher-grade bottleneck genes in TNBC PPI regulatory network include CCL2, CCND1, CDH1, FN1, PLEK, CAT, CFTR, CREBBP, VWF, IL6, DCN, ESR1, IL2RB, PPARG, LRRK2, HPGDS, LIPE, FYN, KDR and GNG11 as per the decreasing order of highest bottleneck score (**Fig. 4B**). Similarly, we also found 20 top betweenness centrality, IL6, FN1, ESR1, CDH1, PTPRC, JUN, CAV1, PPARG, ERBB2, PLEK, CAT, CCND1, EZH2, SOX2, MKI67, FYN, LRRK2, CFTR, VWF and CD34.

**Fig. 4:**
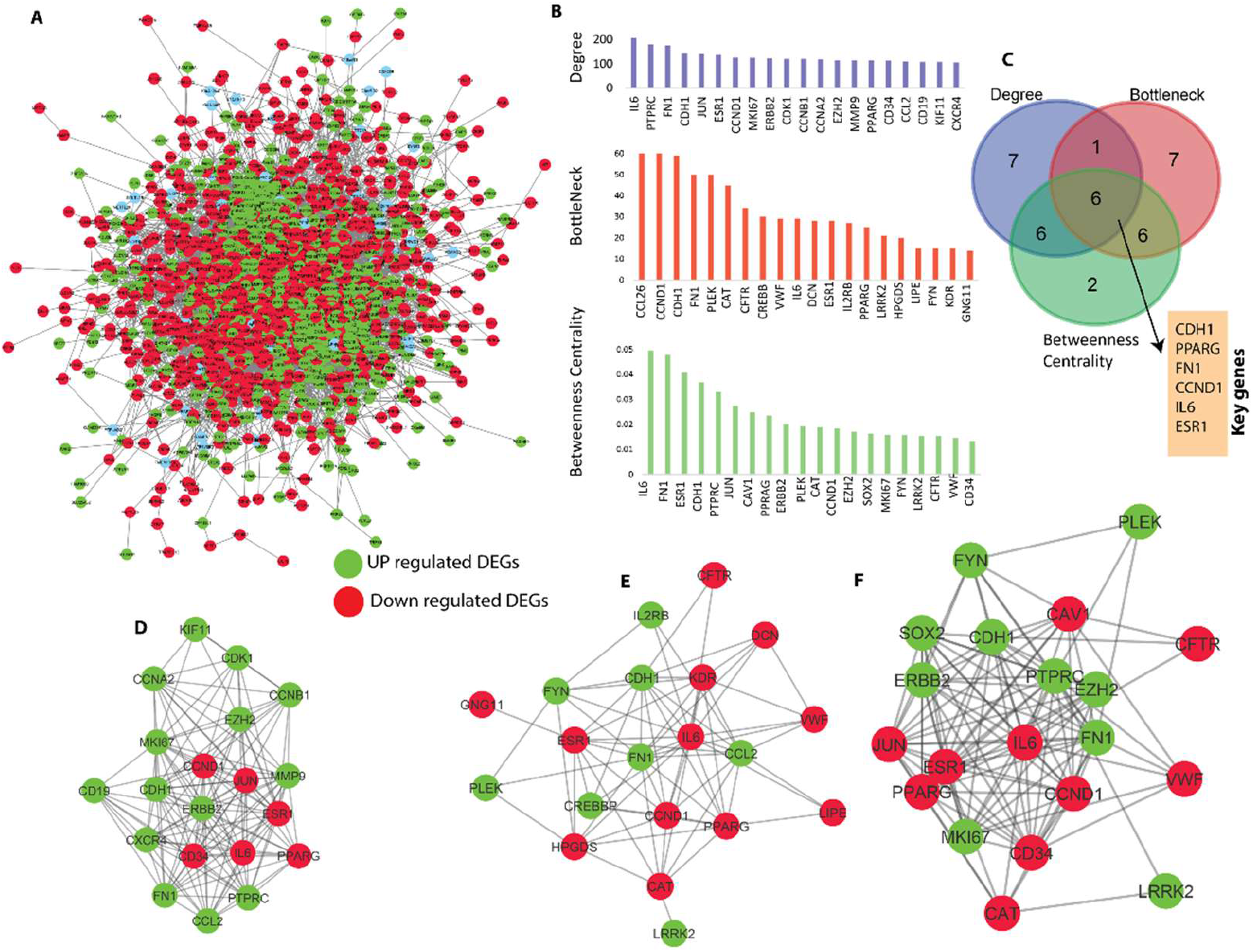
PPI regulatory network of TNBC DEGs. (**A**) PPI network of DEGs. Red colour and green colour nodes show downregulated and upregulated genes of TNBC *vs*. non-TNBC respectively. (**B**) The bar plot shows topological properties (degree, bottleneck and betweenness) (**C**) Venn diagram shows the number of common genes among various centralities of top 20 genes topological parameters i.e., degree, bottleneck and betweenness centrality. (**D-F**) Top hubs, bottleneck and betweenness centrality genes of regulatory network retrieved from PPI network of TNBC.

To identify key genes in the PPI network, top 20 genes from each of the three topological properties were taken to compare with each other. After analysing the topological properties (degree, bottleneck and betweenness centrality), and tracing them to top 20-degree (hub genes) CDH1, PPARG, FN1, CCND1, IL6 and ESR1 were found to be common genes for all topological properties suggesting the significance of these 6 genes in TNBC. The Venn diagram shows common genes of topological properties (**Fig. 4A to 4F**).

### Prediction of mRNA expression pattern of key genes from TCGA

The expression pattern of identified key genes in general breast cancer tissue samples was predicted from mRNA expression using GEPIA [http://gepia.cancer-pku.cn/] online server where p-value is P < 0.01. The expression pattern of CDH1, FN1, CCND1 and ESR1 was found to be significantly up regulated in breast cancer tissue samples as compared to that of non-cancerous tissue samples while IL6 and PPRAG were down regulated (**Fig. 5**). This confirms the differential expression of these genes in other sub types of breast cancer as well.

**Fig. 5:**
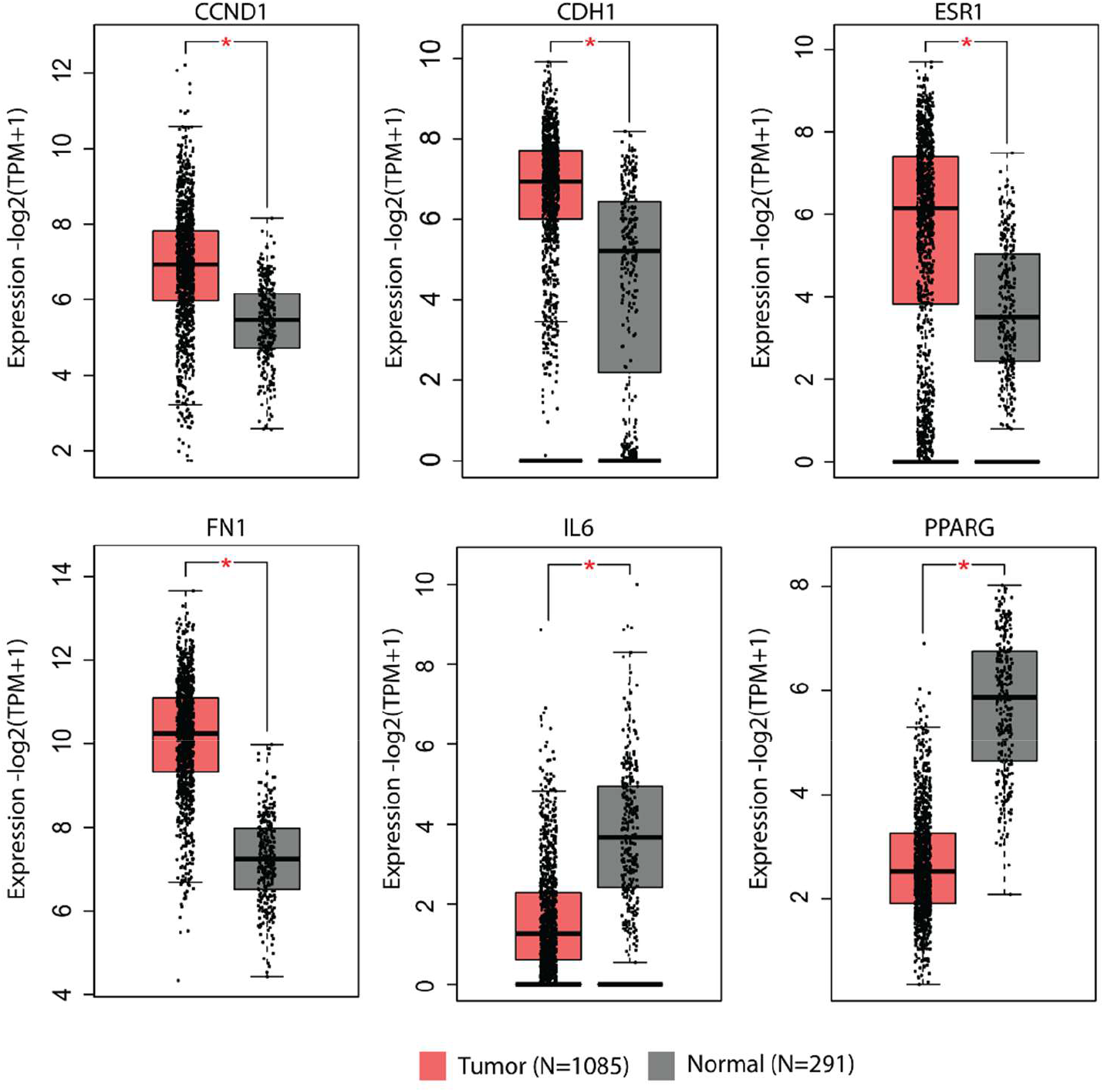
mRNA expression of key genes in TNBC network which compare between non-TNBC (black) and breast cancer patients (red) using GEPIA database: Gene expression levels of CDH1, CCND1, ESR1, FN1, IL6 and PPARG.

The mRNA expression of CCND1, ESR1, IL6 and PPARG genes was found to be related to overall survival. The Kaplan Meier (KM) plot also showed that these genes were significantly associated with breast cancer (**Fig. 6)**. These genes can be correlated with increasing or decreasing risk of TNBC progression; hence, they can be potential prognostics markers or drug targets in breast carcinoma.

**Fig. 6:**
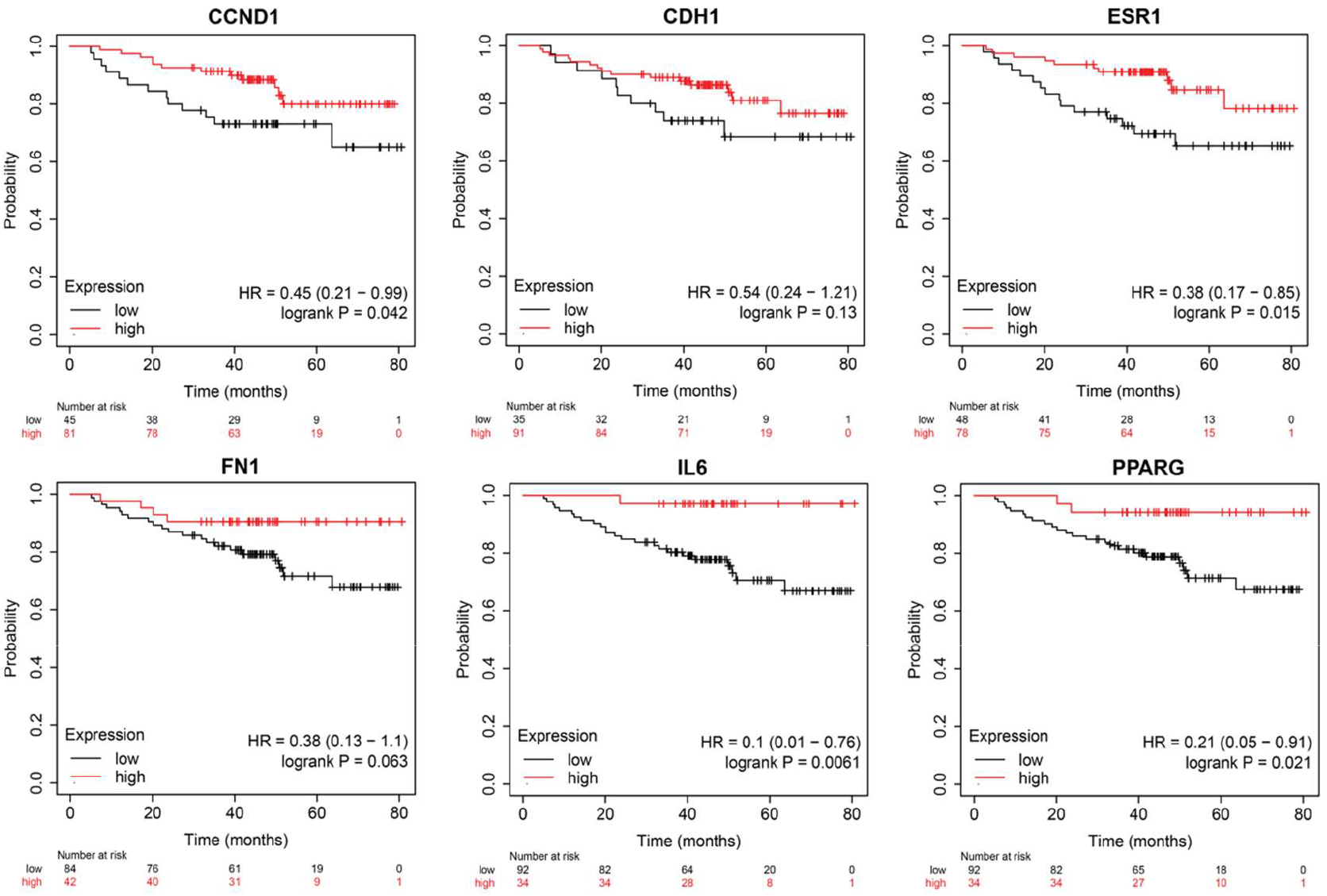
Kaplan–Meier survival curves comparing the high and low expressions of TNBC for six key genes: CCND1, CDH1, ESR1, FN1, IL6 and PPARG.

### Subtype-specific expression patterns of hub genes in breast cancer

To conduct a more in-depth analysis of the expression patterns of the identified hub genes, transcriptomic data from TCGA breast cancer cohorts were evaluated utilizing the UALCAN platform. The expression levels of CCND1, CDH1, ESR1, FN1, IL6, and PPARG were assessed across various molecular subtypes of breast cancer, which include luminal, HER2-positive, and triple-negative breast cancer (TNBC), in addition to normal breast tissues. The analysis uncovered subtype-specific variations in gene expression patterns. Notably, ESR1 expression was significantly diminished in TNBC samples (**Fig. 7A**) when compared to luminal breast cancer, aligning with the lack of estrogen receptor signalling that is the characteristic of TNBC. Likewise, the expression levels of CDH1 and PPARG were found to be lower in TNBC samples in comparison to luminal tumors (**Fig. 7B, 7C**). Conversely, FN1 and IL6 exhibited relatively elevated expression in TNBC compared to luminal breast cancer subtypes (**Fig. 7D, 7E**), indicating their potential involvement in extracellular matrix remodelling and inflammatory signalling pathways linked to aggressive tumor phenotypes. CCND1 expression was detected across several breast cancer subtypes, showing moderate variation among the groups (**Fig. 7F**). These results reinforce the expression patterns associated with the subtypes of the identified hub genes and further emphasize their possible role in the biology of TNBC.

**Fig. 7:**
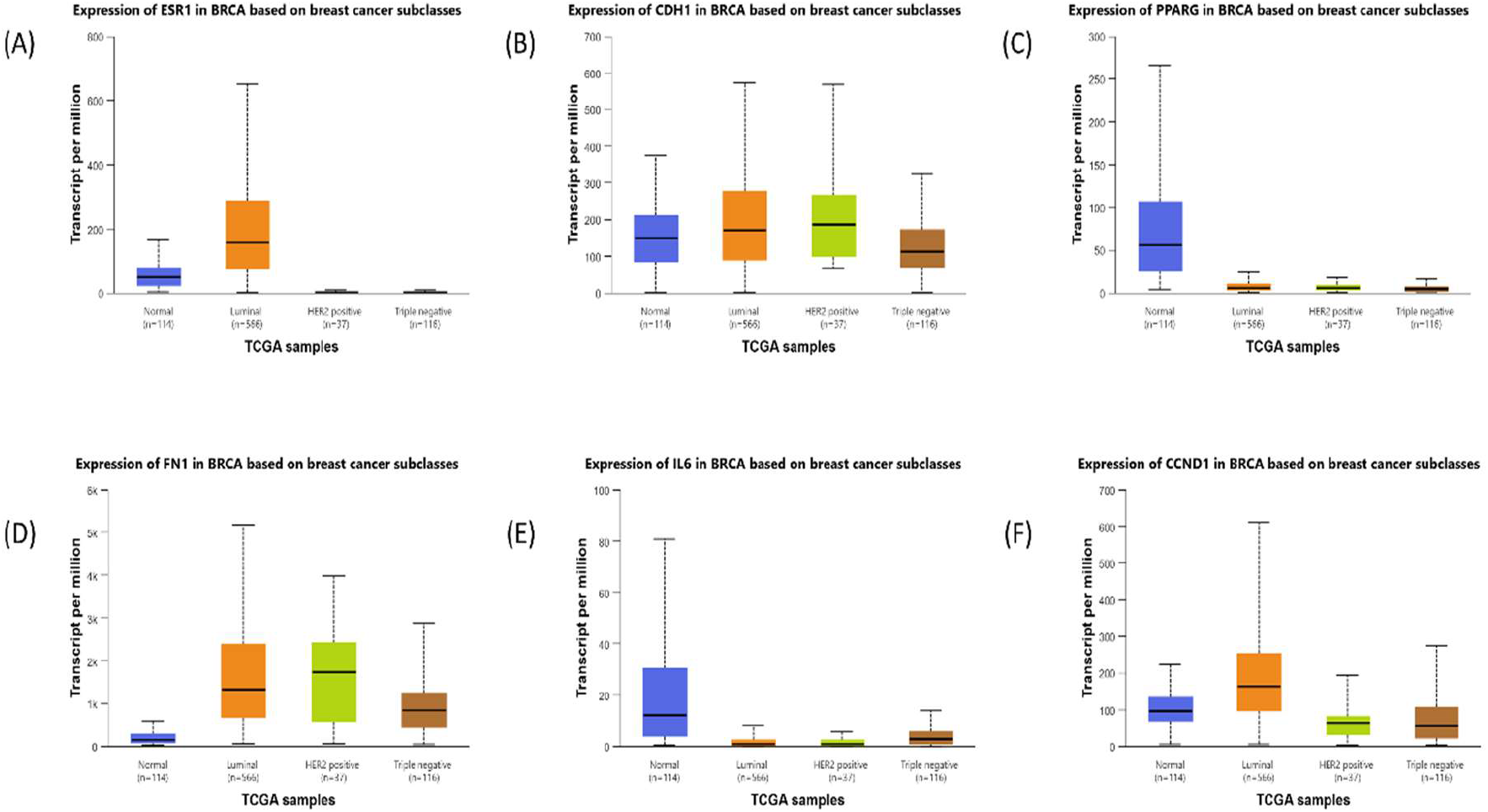
Subtype-specific expression patterns of hub genes in breast cancer

### Experimental validation of hub gene expression in TNBC cell line

The results from the qRT-PCR analysis revealed significant subtype-associated differences in gene expression between the two cell lines (**Fig.8**). Notably, IL6 expression was significantly higher in MDA-MB-231 cells compared to MCF7 cells, indicating an increase in inflammatory signalling within the TNBC model. Likewise, FN1 expression was elevated in MDA-MB-231 cells, suggesting enhanced interaction with the extracellular matrix and potential involvement in the processes of cell migration and invasion that are associated with aggressive breast cancer phenotypes. Conversely, the expression levels of CDH1 and ESR1 were markedly diminished in MDA-MB-231 cells in comparison to MCF7 cells, which aligns with the loss of epithelial characteristics and estrogen receptor signalling that are typically observed in triple-negative breast cancer. CCND1 expression exhibited a moderate increase in MDA-MB-231 cells, indicating possible changes in cell cycle regulation. Interestingly, PPARG expression was found to be elevated in MDA-MB-231 cells relative to MCF7 cells.

**Fig. 8:**
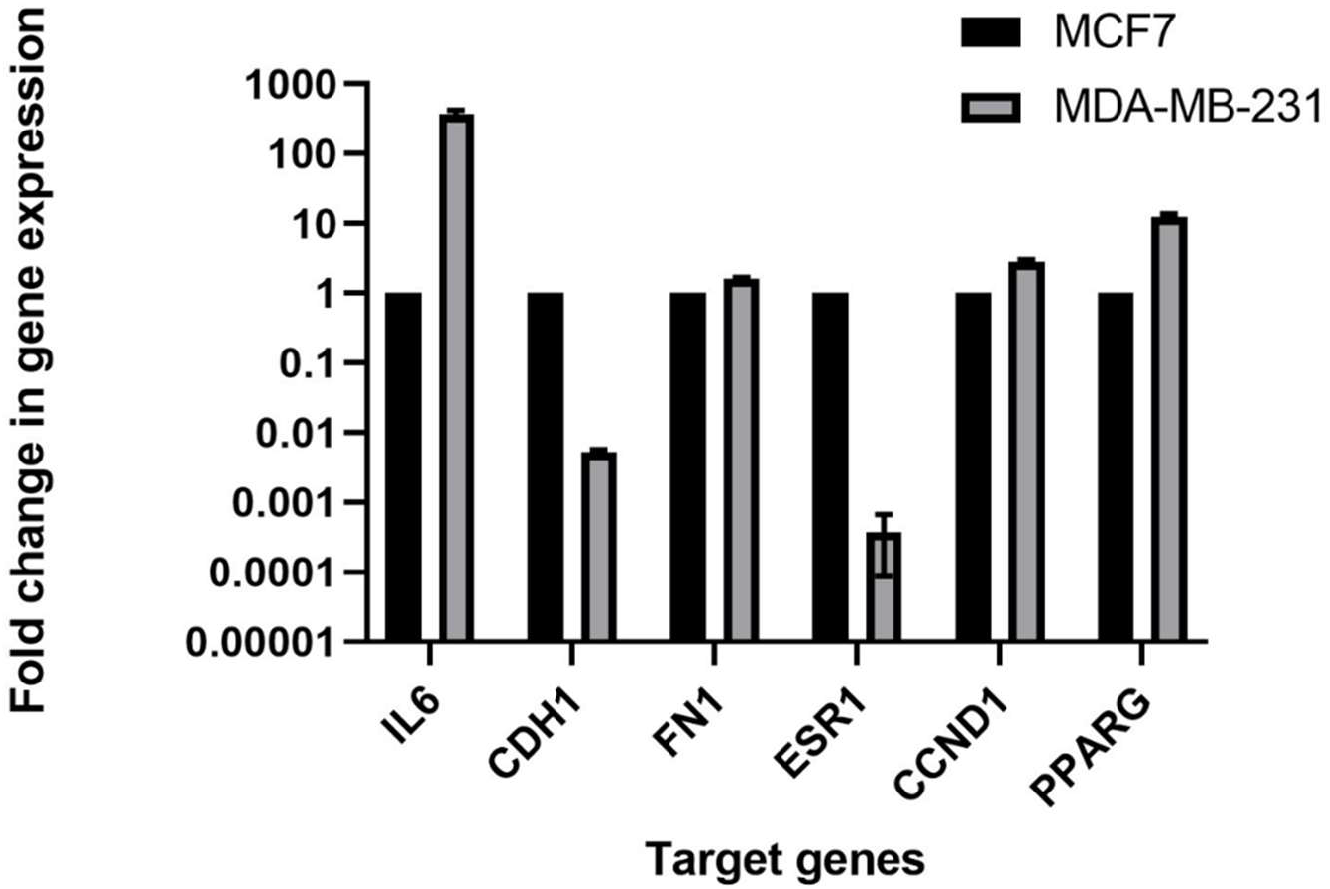
Validation of hub genes expression at transcript level. Total RNA was isolated from TNBC and non-TNBC cells, reverse transcribed and cDNA was amplified by real time PCR using specific primers.

Although this expression pattern diverges from the subtype distribution noted in TCGA datasets, such discrepancies may reflect variations between patient tumor samples and in vitro cell line models, including tumor heterogeneity and the metabolic adaptations that occur in cultured cancer cells.

## Discussion

Network analysis and functional enrichment are well known techniques to find novel potential key genes for any biological network [26-28]. In our study, DEGs between TNBC and non-TNBC tissues of patients in two datasets from GEO were analysed. Microarrays have been commonly used in recent decades to classify DEGs and pathways underlying TNBC pathogenesis. Most microarray data in these studies is predominantly from the PBMC and tissue samples. This study highlights the genes associated with changes in cellular structure and disease progression. Predicting good candidate genes before experimental analysis will save time and effort. Meta-analysis with two microarray datasets from breast tissue samples had shown a total 1810 DEGs. To find the common transcriptomic signatures, meta-analysis was done on human gene expression data of TNBC individuals, and then the string data network was constructed.

Using the network analysis, *CCND1, CDH1, ESR1, FN1, IL6 and PPARG* were identified to be key players involved in TNBC pathogenesis. CCND1 (Cyclin D1) plays a role in cell cycle regulation and has been reported to be overexpressed in various tumors including TNBC tumors. Association of higher expression of CCND1 with aggressive nature of TNBC and poor prognosis was also reported [30].CDH1 (E-cadherin) is another gene identified to be a key regulator in in the maintenance of epithelial tissue integrity. It is involved in the regulation of cell-cell adhesion, cell-matrix interactions and is responsible for mediating calcium-dependent interactions between adjacent cells. Loss of CDH1 expression is associated with various cancers, including breast, gastric, and ovarian cancer [31]. It is used as a biomarker to classify the subtype and also determines the patient outcome. ESR1 is a well-established therapeutic target in hormone receptor-positive breast cancer, where it is targeted with hormone therapy such as tamoxifen or aromatase inhibitors (4). FN1 is being investigated as a therapeutic target in breast cancer, as inhibition of its expression or function may reduce cancer cell migration and invasion [32, 33]. IL-6 has been shown to promote breast cancer cell proliferation and survival; targeting IL-6 pathway with drugs such as tocilizumab or siltuximab is under investigation as a potential therapy to treat breast carcinoma [34, 35]. PPARG is also being investigated as a therapeutic target in breast cancer; the inhibition of its expression or function may reduce cancer cell proliferation and promotes survival [36, 37]. Drugs that target PPARG, such as pioglitazone or GW9662, are also under investigation as potential therapies in breast cancer[38]. Thus, these genes can be used as biomarkers for diagnosis and prognosis. However, using them alone as reported in previous studies may not provide a complete picture of the patient’s disease status.

Considering the importance of the identified genes in TNBC, this study proposes the use of these markers in in combination with other clinical, pathological and imaging data for a patient specific treatment regimen. Additionally, the cut-off levels for these biomarkers to be considered as positive or negative may vary between studies and should be considered accordingly. These biomarkers are not only being investigated in breast cancer but also in other cancers as they may have different implications depending on the type of cancer and the stage of the disease. Further research is needed to fully understand their implications in cancer and their potential use in clinical practice.

## Conclusion

Current study identified multiple important genes, which showed differential expression in TNBC patients compared to control through DEG analysis and network building. Our integrated analysis suggested that the identified genes could be potential diagnostic and prognostic biomarkers of TNBC. However, as the current findings were extracted by applying computational biological approach, and preliminary experimental work in breast cancer cell lines, efficient experimental research is required to fully understand the underlying molecular mechanisms regulated by the identified key genes in TNBC.

## Authors Contributions

Alok Kumar: Performed the bioinformatic studies and analysis, written the first draft, Mohammad Kashif, Md. Zubbair Malik: Performed the analysis, Gauri Shankar Upadhyay: Performed gene validation studies *in vitro*, Naidu Subbrao: Conceptualization, Maitreyi S Rajala: conceptualization, writing, review and editing,

## Ethics approval

Not applicable

